# Convergent effects of neurodevelopmental disorder-associated variants at mitochondria

**DOI:** 10.64898/2026.02.26.708316

**Authors:** Maxine I. Robinette, Jada B. Gundy, Xinyan Leng, Duc Duong, Ananth Shantaraman, Liang Shi, Elizabeth A. Candelario, Anson Sing, Shubhangi Vibhor Garg, Weibo Niu, Nicholas T. Seyfried, Steven A. Sloan, Zhexing Wen, Joseph F. Cubells, Erica Duncan, Jennifer G. Mulle, Victor Faundez, Gary J. Bassell, Ryan H. Purcell

## Abstract

Investigations into the molecular pathogenesis of clinically defined neurodevelopmental disorders (NDDs) including autism spectrum disorders (ASD) and schizophrenia (SCZ) have produced evidence implicating dysfunctional mitochondrial metabolism. However, the functional connection between risk variants and mitochondrial proteins largely remains unclear. We tested the hypothesis that proteins encoded by NDD-associated copy number variants (CNVs) and SCZ risk genes are enriched within the mitochondrial interactome. We found that NDD- and SCZ-associated genes exhibit mitochondrial association comparable to their overlap with synaptic proteins, with interaction networks converging most strongly on mitochondrial translation. Two high-risk CNVs, the 3q29 deletion (3q29Del) and the 22q11.2 deletion (22q11Del), confer similar risks for ASD and SCZ and have independently been linked to mitochondrial phenotypes. To test whether these CNVs produce convergent effects on mitochondrial proteins in developing human neural tissue, we generated an isogenic series of 3q29Del and 22q11Del induced-pluripotent stem cells (iPSCs) and differentiated them into forebrain cortical organoids. Quantitative proteomic analysis showed high similarity in the profiles of dysregulated proteins in 3q29Del and 22q11Del compared to isogenic controls. Enrichment analysis of proteins altered in both variants revealed significant convergence on the mitochondrial ribosome and translation machinery. Furthermore, manipulation of mitochondrial translation elicited similar proteomic and functional responses in organoids and neural progenitor cells across both CNVs. These findings indicate that NDD-associated genes have rich interactions with mitochondrial proteins and that two of the strongest risk factors for NDDs may similarly disrupt neural mitochondrial metabolism through impaired mitochondrial translation.

## Introduction

Recurrent copy number variants (CNVs), defined as deletions or duplications of genomic segments, confer substantial risk for neurodevelopmental disorders and are often associated with early disease onset and increased clinical severity [1–3]. Because CNVs are oligo- or polygenic and can perturb multiple genes simultaneously, they also serve as a model for dissecting pathway-level mechanisms underlying polygenic disease [4, 5]. However, a central challenge remains in determining molecular and cellular mechanisms for how specific CNVs perturb the developmental trajectory of the human central nervous system in ways that increase liability for NDDs.

There is a paradox inherent in the current landscape of NDD genetics: many rare variants of high effect are associated with NDDs [6, 7], yet these distinct variants produce phenotypes that converge at the clinical level onto robustly characteristic diagnostic entities. These observations introduce a fundamental question in the neurobiology of NDD genetics as to whether each high-risk variant produces a largely unique molecular disruption of neurodevelopment, or whether distinct variants converge on shared pathway signatures that could support mechanism-based therapeutic strategies. While psychiatric genomics has been highly successful in mapping risk loci, linking these loci to convergent cellular processes remains challenging.

Extensive functional studies, conducted in parallel with genetic investigations, have repeatedly implicated mitochondrial dysfunction in NDD pathophysiology. Studies in mouse and human models, such as post-mortem brain samples and induced pluripotent stem cells (iPSCs), have frequently revealed disrupted mitochondrial morphology, altered ATP production, and changes in metabolic flux as core features across a range of disorders [8–16]. Yet, these functional data exists in apparent tension with genomic findings, which largely implicate neuronal development and synaptic gene annotations [6, 17]. The functional impact of mitochondrial annotated variants [18, 19] has remained uncertain unless mitochondrial annotated genes from population-scale genetic studies were to be high connectivity nodes within the mitochondrial proteome [20]. Here, we tested this model focusing on two CNV with strong NDD phenotypic overlap, 22q11.2 and 3q29 deletion syndromes.

3q29 deletion (3q29Del) and 22q11.2 deletion (22q11Del) are two recurrent CNVs that confer the highest known lifetime risk for developing SCZ [7, 21–23] and are associated with high risk for ASD [24]. Despite impacting entirely distinct genomic loci, both CNVs produce somewhat similar neurodevelopmental outcomes, including feeding difficulties [25–27], palatal abnormalities [26, 28], motor delays [25, 26], language delays [25, 29], mean full-scale IQ in the low 70s [26, 30], executive function deficits [30, 31], and increased rates of several psychiatric syndromes including attention deficit-hyperactivity disorder (ADHD) [30, 32], SCZ and other psychotic disorders, and generalized anxiety disorder [30, 33]. Together, this notable convergence in neurodevelopmental phenotypes, despite non-overlapping genetic lesions, suggests that 3q29Del and 22q11Del may disrupt shared neurodevelopmental pathways and provides a strong rationale for directly comparing their molecular consequences in human neural models.

The 22q11Del (prevalence ∼1:2,000-4,000) confers a 25-30% lifetime risk for psychosis and typically results in a 3.0Mb deletion encompassing approximately four dozen protein-coding genes, several of which localize to mitochondria [34–38]. In contrast, the 3q29Del (prevalence ∼1:30,000) results in hemizygosity of 22 protein-coding genes yet is associated with an even higher approximate psychosis burden of ∼40% and contains a single annotated mitochondrial-localizing protein [39–42]. Both 3q29Del and 22q11Del are associated with mitochondrial dysfunction, particularly affecting oxidative phosphorylation (OxPhos). Specifically, an analysis in mouse and human 3q29Del models implicated impaired metabolic flexibility in the glycolysis to OxPhos transition of developing neurons [43]. Studies of 22q11Del showed reduced activity in the electron transport chain (ETC) and ATP deficits [44], and a compensatory shift towards glycolytic metabolism in plasma from carriers [45]. Further dissection of potential driver genes has implicated *SLC25A1* and *MRPL40*, a component of the mitochondrial ribosome, in altered mitochondrial translation [46–48]. These mitochondrial impairments in 22q11Del have been shown to impact cortical circuitry in a mouse model [16]. Still, the mechanistic connection from CNV oligogenic haploinsufficiency to neuronal mitochondrial dysfunction is unclear.

Here, we sought to isolate the impact of 3q29Del and 22q11Del in an isogenic background to reduce variability and identify primary molecular consequences in developing human neural tissue. We engineered isogenic human iPSC lines harboring either the 3q29Del or the 22q11Del and differentiated them into forebrain cortical organoids to model early human corticogenesis. Using quantitative proteomics across neurogenic and gliogenic stages, we find that both CNVs induce highly similar proteomic disruptions. These changes converge strongly on mitochondrial pathways, most prominently constituents of the mitochondrial ribosome and other components of the mitochondrial translation machinery. We then used metabolic and pharmacologic perturbations to challenge mitochondrial gene expression and translation as a test of a shared functional vulnerability linking genetically distinct NDD risk variants.

## Methods

### Cell Line Engineering

Isogenic 22q11.2 deletion (22q11Del) iPS cell lines were generated by the SCORE method [49] from the same neurotypical control parent line as reported previously [43]. Low copy repeat (LCR) segments A (hg38 chr22:18712538-18878770) and D (chr22:21515447-21679525) of the 22q11Del locus were searched for unique gRNA sequences using CHOPCHOP (https://chopchop.cbu.uib.no) [50]. BLAST identified ∼44kb of 99% identical sequence and candidate gRNA sequences were checked against reference sequence for LCRs B and C to avoid alternative variants.

gRNA sequences were cloned into pSpCas9(BB)-2A-Puro (PX459) V2.0, which was a gift from Feng Zhang (Addgene plasmid #62988; http://n2t.net/addgene:62988; RRID: Addgene_62988) [51], reverse transfected into iPS cells, and screened for double-strand break efficiency (GeneArt gDNA cleavage, Thermo). Single-cell clones transfected with most efficient gRNA (“gRNA_2”, GCTGTAGTATCATCAATCAC, targeting hg38 chr22:18820811 & chr22:21255283) were briefly selected for episomal plasmid conferred puromycin resistance, expanded, and isolated by serial dilution. 100 clones were screened for alteration in copy number of *PRODH* by TaqMan gDNA copy number assay (assay ID: Hs04080831_cn, referenced to *RNAseP* Thermo 4401631) and four appeared to be hemizygous. All four clones were subsequently tested (Supp. Fig.1) for copy number of *TUBA8* (Hs04094671_cn, outside, centromeric), *HIC2* (Hs00765260_cn, outside, telomeric), *DGCR8* (Hs01290445_cn, deletion between LCRA-B), *KLHL22* (Hs04092938_cn, deletion between LCRB-C), and *SNAP29* (Hs01130938_cn, deletion between LCRC-D).

**Figure 1.**
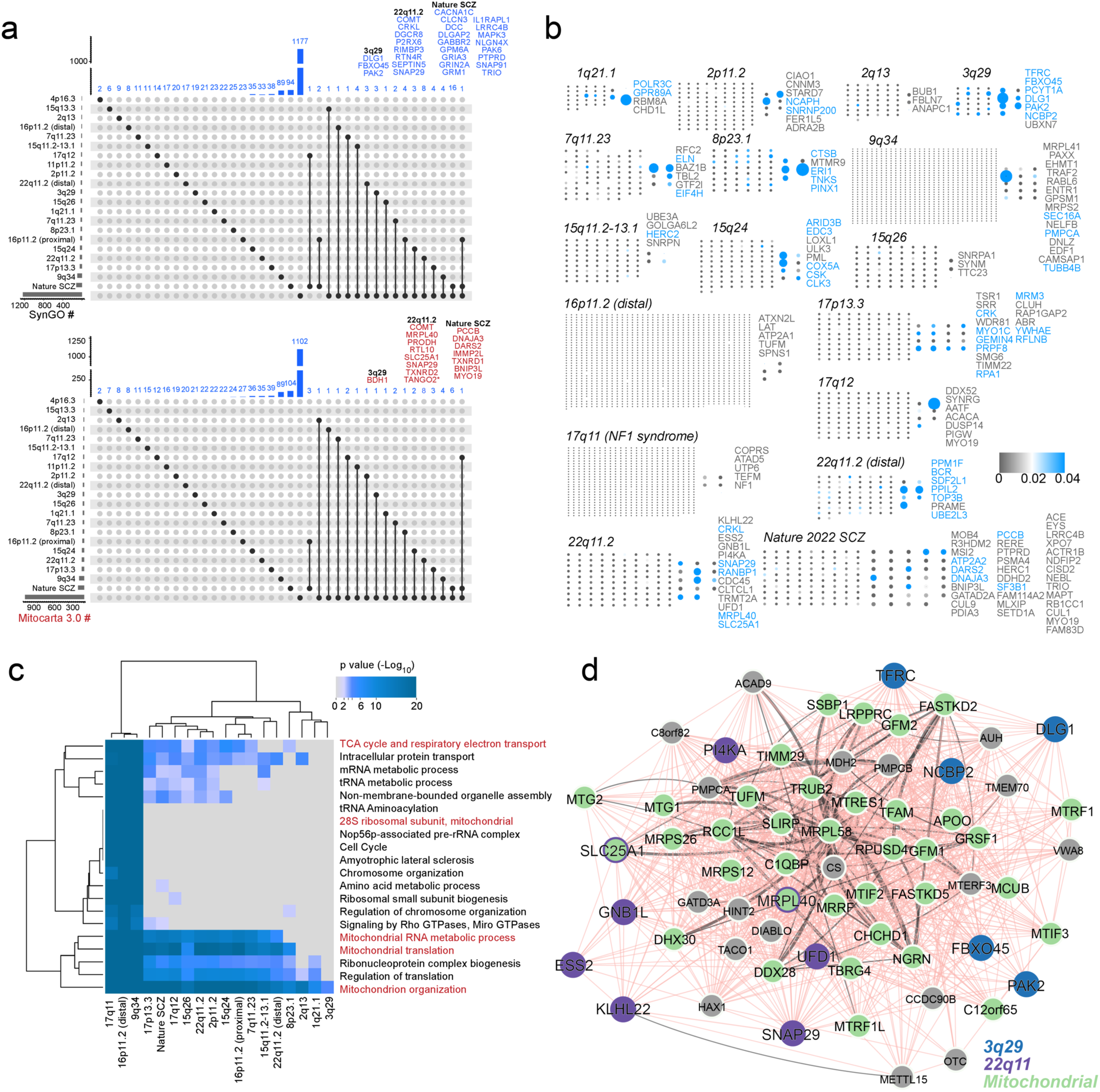
Neurodevelopmental Disorder (NDD)-Associated CNVs and SCZ Risk Genes Form Nodes of High Connectivity within a Mitochondrial Interactome. (a) Synaptic and mitochondrial proteins are similarly represented among Neurodevelopmental disorder (NDD)-associated CNVs and SCZ risk genes. UpsetR plots depicts the intersection of (NDD)-associated CNVs and SCZ risk genes with the synaptic (SynGO) and mitochondrial (MitoCarta 3.0) annotated proteins. The proportion of either synaptic or mitochondrial annotated genes represented among either CNVs or SCZ risk genes is the same. Two tailed *Fisher Exact Probability Test p=* 0.133 and 0.0769 respectively (b) Neurodevelopmental disorder (NDD)-associated CNVs and SCZ risk genes form nodes of high connectivity in a mitochondrial interactome. All nodes are proteins present in the mitochondrial interactome. Nodes grouped to the left represent genes from the mitochondrial interactome residing outside the CNV or SCZ gene list. Nodes clustered to the right depict genes belonging to the mitochondrial interactome and residing in the indicated CNV or SCZ gene list. Edges have been removed for clarity. Heat map depicts node connectivity measured by betweenness centrality. Blue nodes are nodes of high connectivity. Node size is proportional to betweenness centrality score. (c) The interactomes of NDD CNV and SCZ risk proteins strongly implicates mitochondrial ontologies preferentially annotated to mitochondrial protein synthesis terms. (d) Illustration of model of NDD-associated variants across the genome converging on mitochondria annotated genes based on GeneMania protein-protein interactions (black) and genetic interactions (salmon).

### Cell culture and cortical organoid differentiation

iPSC lines grew to 80-90% confluency in hESC-qualified Matrigel-coated plates and were maintained with mTeSR+ medium (STEMCELL Technologies). Dorsal forebrain patterned cortical organoids were generated based on a published protocol [52]. On day 0 of in vitro differentiation (DIV0), colonies were washed with sterile PBS and incubated with Dispase (0.35mg/mL) until detachment. Detached colonies were transferred to a 15-mL conical tube using pre-warmed DMEM/F12 and a pre-wetted 25-mL serological pipette, rinsed three times with warm DMEM/F12, and resuspended in mTeSR+ supplemented with 10 µM Y-27632. Aggregates were then transferred to 10-cm plates that had been pre-rinsed with Anti-Adherence Solution (STEMCELL Technologies) and washed twice with DPBS.

After 48 hours, developing neural spheroids were transitioned to daily neural induction media (Dulbecco’s Modified Eagle Medium/Nutrient Mixture F-12 (DMEM/F12) with HEPES, 20% knockout serum replacement, 1% non-essential amino acids, 0.5% GlutaMax, 1% penicillin/streptomycin) freshly supplemented with 5uM of Dorsomorphin and 10uM of SB-431542 for DIV2-6. From DIV7–16, neural induction media was replaced with neural differentiation medium (Neurobasal-A, 2% B27 supplement without vitamin A, 1% GlutaMAX, and 1% penicillin–streptomycin) with daily addition of 20 ng/mL epidermal growth factor (EGF) and 20 ng/mL fibroblast growth factor-2 (FGF2). From DIV17–25, organoids were fed with the same medium containing EGF and FGF2 every other day. On DIV26, the medium was supplemented with 20 ng/mL brain-derived neurotrophic factor (BDNF) and 20 ng/mL neurotrophin-3 (NT-3), and organoids were maintained with feeding every other day. From DIV 43 onward, organoids were fed twice weekly with unsupplemented neural differentiation medium.

For microplate viability assays, iPS cells were differentiated to neural progenitors by one of two methods. Cells were either differentiated by standard dual-SMAD inhibition as described previously [43] or by a recently reported method to improved synchronization of neurogenesis [53]. Briefly, homogeneous iPSC cultures were differentiated in high-density monolayers in the presence of 2uM XAV939 (day 0-3), 100nM LDN193189 (day 0-10), and 10μM SB431542 (day 0-10) in Essential 6 medium (Gibco). Cells were further matured for 10 days in N2/B27 medium (1:1 NB:DMEM/F12 basal medium supplemented with 1xN2 and B27 minus vitamin A) before passaging into assay plates or vials for cryostorage. Neural progenitor cell and organoid differentiations were performed by multiple experimenters at both Emory University School of Medicine and the Fralin Biomedical Research Institute at Virginia Tech with no discernable differences by laboratory.

### Immunofluorescence

Organoids were fixed in 4% paraformaldehyde/1x PBS and dehydrated in 30% sucrose for 72 hours. Organoids were then marked with one drop of methylene blue, embedded (Tissue-Tek OCT Compound, Sakura, Reference number 4583), and frozen at –80 °C until sectioning. Blocks were sectioned at 14μm and melted onto charged slides (Premiere Microscope Slides ½ Gross). Slides were stored at -80°C until staining. Mounted organoid sections were washed in 1x PBS for 10 minutes. Sectioned organoids were then washed in 1x PBS, blocked (5% Normal Donkey Serum, 0.1% Triton X-100, 5% BSA, 1xPBS) and incubated with primary antibodies (SOX2 goat R&D AF2018 1:40, NES mouse Abcam ab22035 1:500, MAP2 guinea pig Synaptic Systems 188 004 1:500) in blocking solutions overnight at 4°C in a humidified chamber. Samples were then washed and incubated with secondary antibodies and DAPI (Biotium 40043, 1:2000) in blocking solution for 1 hour at RT. Slides were washed and cover glasses were mounted with Vectashield Vibrance Antifade Mounting Medium (ref. H-1700-10) and stored away from light prior to imaging.

### Tandem Mass Tag Proteome Preparation

At DIV 50, 100, and 150, neural media was aspirated and organoids were washed once in DPBS. Organoids were triturated in 150uL of 8M Urea buffer (in 100mM Tris-HCl pH 8.0 with 1x HALT Protease and Phosphatase inhibitor) 20x. Samples were vortexed for 10 sec, sonicated 3x 5 seconds at 30% power, and stored at -80C. Prior to processing, samples were further homogenized in Rino bead tubes and as detailed previously [54].

### Weighted Co-Expression Network Analysis

Normalized TMT proteome data were analyzed with a modified version of the bioinformatic pipeline for weighted co-expression network analysis (WGCNA) [55, 56]. We generated a signed network, soft power (beta) = 17, using biweight midcorrelation. Function mergeCutHeight was set to 0.07, deepSplit=2, and minimum module size was 20. Module membership was determined by signed kME, the correlation with first principal component (module eigenprotein) of each module.

### Proteome Analyses

Differential expression (DEx) analysis for TMT data was performed by one-way ANOVA using the parANOVA.dex script (Github: edammer) [54]. Overlap analysis was conducted in the *GeneOverlap* package in R [57] and fold enrichment was determined by hypergeometric test. Spearman correlations were performed in R and plotted in ggplot. Venn diagrams were plotted using the R package *VennDiagram* [58]. The percent of expected DEx mitochondrial proteins (Fig. 3h) was determined by calculating the percent of Mitocarta3.0 proteins [59] that were detected in the dataset (1216 of 8194 proteins = 14.84%). We then estimated that for 194 DEx proteins (midpoint of 3q29Del and 22q11Del DEx protein numbers), we would expect 29 to be mitochondrial. Heatmaps were generated with the R package *pheatmap* [60]. DEx due to galactose treatment was determined in each genotype by FDR-adjusted Q<0.05 and absolute value of log_2_-fold change > 0.1.

### Single-cell RNA-seq

DIV 50 dorsal cortical organoids were differentiated as described above and dissociated using a papain-DNase method based on a published protocol [61] and as described previously [43]. 10x Genomics 3’ v3 chemistry, *CellRanger*, and *Seurat* v5 were applied as detailed previously [43]. Doublets were identified and excluded using *DoubletFinder* [62]. Transcriptomically defined clusters were combined based on canonical marker gene expression into seven cell types. Cell type proportions were analyzed using the *propeller() function of speckle* [63].

### Bulk RNAseq

Whole RNA was isolated from QIAshredder-lysed organoids by a phenol-chloroform and column method (miRNeasy kit, Qiagen). Messenger RNA (mRNA) isolation, library preparation, sequencing, and read count quantification were performed by Novogene. Briefly, mRNA was purified from total RNA using poly-T oligo-attached magnetic beads, fragmented, and cDNA was synthesized from the first strand using random hexamer oligos followed by the second strand cDNA synthesis. After library check and QC, libraries were pooled and sequenced on an Illumina platform. Raw reads (44.8M +/- 4.1M) of fastq format were processed, trimmed, and low-quality reads were removed.

Reads were mapped to human hg38 by HISAT2 (2.2.1) and reads counts per gene were determined by featureCounts (2.0.6). FPKM (Fragments Per Kilobase of transcript sequence per Millions base pairs sequenced) of each gene was calculated based on the length of the gene and reads count mapped to this gene.

Differential expression analysis was performed in DESeq2 [64]. All genes with total counts across all samples > 500 were kept for analysis. Differential expression was determined by treatment (glucose vs. galactose) within each genotype (Control, 3q29Del, 22q11Del) at FDR-corrected P<0.05.

### Immunoblotting

Organoids were lysed in ice cold lysis buffer (150mM NaCl, 25mM Hepes, 10mM MgCl2, 1mM EDTA, pH 7.4) supplemented with protease/phosphatase inhibitors (HALT, Thermo) and 1% Triton X-100 (Sigma). Samples were mechanically homogenized using a BeadBug homogenizer and then processed through QIAshredder spin columns to clear large debris. Lysates were sonicated, incubated on ice for 30 minutes and centrifuged at 15,000 × g for 10 minutes at 4 °C to further remove insoluble cell debris. Supernatant was collected, and protein concentrations were determined via BCA assay. Protein lysates were prepared in 4x loading buffer (BioRad) and heat-denatured at 95°C for 5 minutes, then either stored at -80°C or were resolved on a gel the same day. 20ug of protein was electrophoresed on a 4-20% mini-PROTEAN TGX precast protein gels for one hour. Gels were transferred (Trans-Blot Turbo Transfer System) to a 0.44 μm nitrocellulose membrane and blocked in Intercept (TBS) buffer (LICOR) for 1 hour at RT. Primary antibodies were diluted with equal parts of blocking buffer (1:1) and 1% TBS-Tween 20 and incubated on membranes overnight in a 4°C rocker. The following day, secondaries were added for 1 hour at room temperature. Then, membranes were imaged on a BioRad ChemiDoc Imaging System and normalized with the BioRad Image Lab Software.

### Neural progenitor cell viability

The antibiotic quinupristin/dalfopristin (Q/D) powerfully inhibits mitochondrial translation [65]. Cellular metabolic activity was quantified using a resazurin-based reduction assay. NPCs were seeded in clear-bottom 96-well plates at 50,000 cells per well treated and with Q/D at final concentrations of 1 µM or 5 µM for 72 hours. Q/D stock solutions were prepared in DMSO, and vehicle matched controls were included in the assay (≤0.05% DMSO). Following treatment, resazurin reagent (R&D #AR002) was added directly to the existing culture medium to achieve a final concentration of 10 µg/mL. Plates were incubated for 5 hours at 37 °C in a humidified 5% CO₂ incubator. Fluorescence intensity was measured using a microplate reader with excitation at 450 nm. Background fluorescence from wells containing media and resazurin but no cells was subtracted from all measurements.

### Statistics

Statistical analyses were performed in R version 4.4.1 (2024-06-14) and Prism v10.4.1 (GraphPad). Mitochondrial interactomes were built using the Antonicka et al datasets [66]. Paired interaction nodes and edges were imported into Cytoscape 3.10.2. Node connectivity was determined by betweenness centrality.

## Results

### NDD CNV and SCZ risk genes converge on mitochondrial interactome

We reasoned that mechanistically consequential pathways affected in neurodevelopmental disorders (NDDs) should be recurrent across multiple genetic lesions and also be represented among schizophrenia (SCZ) risk genes that are causal for or associated with NDD. To test this hypothesis, we analyzed 20 recurrent copy number variants (CNVs) and 116 SCZ risk genes curated by the Psychiatric Genomics Consortium and performed an in-silico comparison of the representation of synaptically annotated genes versus mitochondrially annotated genes within these gene sets.

We used the synapse as a well-established mechanistic benchmark for gene enrichment across NDDs [17]. The overall proportion of synaptic versus mitochondrial genes did not differ significantly when considering either all CNVs or all SCZ risk genes (two-tailed Fisher’s exact test, p = 0.133 and p = 0.0769, respectively). Nonetheless, 19 of the 20 CNVs contained SynGO-annotated synaptic genes. The greatest number of synaptic genes was encoded by the 22q11.2 CNV (8 genes; Fig. 1a; 2.9-fold enrichment, p < 0.006). This degree of enrichment was comparable to that observed among the 116 SCZ risk genes, in which 16 SynGO-annotated synaptic genes were identified (Fig. 1a; 2.3-fold enrichment, p < 0.002).

Similarly, Mitocarta 3.0–annotated mitochondrial genes were present in 18 of the 20 CNVs. The 22q11.2 CNV again showed the strongest signal, encoding eight nuclear-encoded mitochondrial genes (Fig. 1a; 3.32-fold enrichment, p < 0.006). In contrast, only seven nuclear-encoded mitochondrial genes were identified among the 116 SCZ risk genes, which did not constitute a significant enrichment (Fig. 1a; 1.1-fold enrichment, p = 0.459). Despite this difference, the pervasive presence of mitochondrial genes across CNVs and SCZ risk loci suggests that the mitochondrion represents a convergence hub across genetic defects and risk factors associated with NDD.

To further assess functional connectivity, we applied an orthogonal strategy that mapped CNV genes and SCZ risk genes onto a high-resolution human mitochondrial BioID proximity interactome, and quantitatively assessed their network properties using graph theory approaches (Fig. 1b). Genes from 16 CNVs were represented as nodes within the mitochondrial interactome, with several exhibiting high connectivity as measured by betweenness centrality (Fig. 1b, blue nodes).

Two salient patterns emerged. First, large CNVs such as 22q11.2 formed extensive mitochondrial interaction networks: 13 genes from the 22q11.2 locus connected to 113 proteins within the mitochondrial interactome. Five of these genes functioned as high-connectivity nodes, including MRPL40 and SLC25A1, which we have previously validated genetically and biochemically to affect mitochondrial translation [48, 67, 68]. Second, the 3q29 CNV displayed a smaller mitochondrial interactome footprint (31 interacting proteins) but the highest proportional enrichment of high-connectivity nodes, with six such genes, including PAK2, which although not annotated in Mitocarta 3.0, we have shown is required for mitochondrial metabolic flexibility [43].

We then subjected all CNV- and SCZ-associated genes present within mitochondrial interactomes to comparative ontological analysis using Metascape. This analysis revealed significant enrichment of pathways related to mitochondrial translation and mitochondrial protein synthesis across both CNV-defined and SCZ-defined interactomes. Protein–protein interaction networks constructed using Antonicka et al. BioID-identified proteins and GeneMania physical and genetic interaction analysis suggest that 22q11Del and 3q29Del proteins separately converge on mitochondrial protein networks (Fig. 1d). Collectively, these in silico analyses support the hypothesis that mitochondrial translation constitutes a shared mechanistic hub linking NDD-associated CNVs and SCZ risk genes.

### Network analysis of 3q29Del and 22q11Del reveals impact on mitochondrial module

We empirically tested the hypothesis that mitochondria and in particular mitochondrial translation are common hubs among NDD associated CNVs. We focused on two CNVs strongly associated with SCZ risk, 22q11Del and 3q29Del, and that displayed high connectivity to the mitochondrial interactome. To this end we engineered the ∼2.5Mb 22q11.2 deletion (22q11Del) into a control iPSC line in which we had previously generated isogenic 3q29Del lines [43]. Four clonal lines were identified and thoroughly characterized (Supp. fig. 1). We differentiated this isogenic cohort of Control (3 clones of parent line), 3q29Del (3 isogenic clones), and 22q11Del (4 isogenic clones) cell lines to forebrain dorsal cortical organoids as described above for up to 150 days in vitro (DIV).

We unbiasedly assessed the impact of each CNV on protein expression during cortical development by performing quantitative tandem mass tag (TMT) mass spectrometry on 45 organoid samples evenly divided by three time points and genotypes (Fig. 2a, d, DIV 50: N=5/genotype; DIV 100: N=5/genotype; DIV 150: N=5/genotype). Normalized expression values for 8,243 proteins were analyzed by weighted co-expression network analysis (WGCNA) and 42 co-expression modules were identified in addition to M0:grey for weakly-connected protein nodes [55] (Fig. 2b). 3q29Del (11 proteins analyzed) and 22q11Del (22 proteins analyzed) CNV-encoded proteins were distributed throughout the network (Fig. 2c). Notably, one protein from each CNV locus was found to be a hub protein of its module (kME > 0.9): MRPL40 (22q11) and NCBP2 (3q29, Fig. 2c).

**Figure 2.**
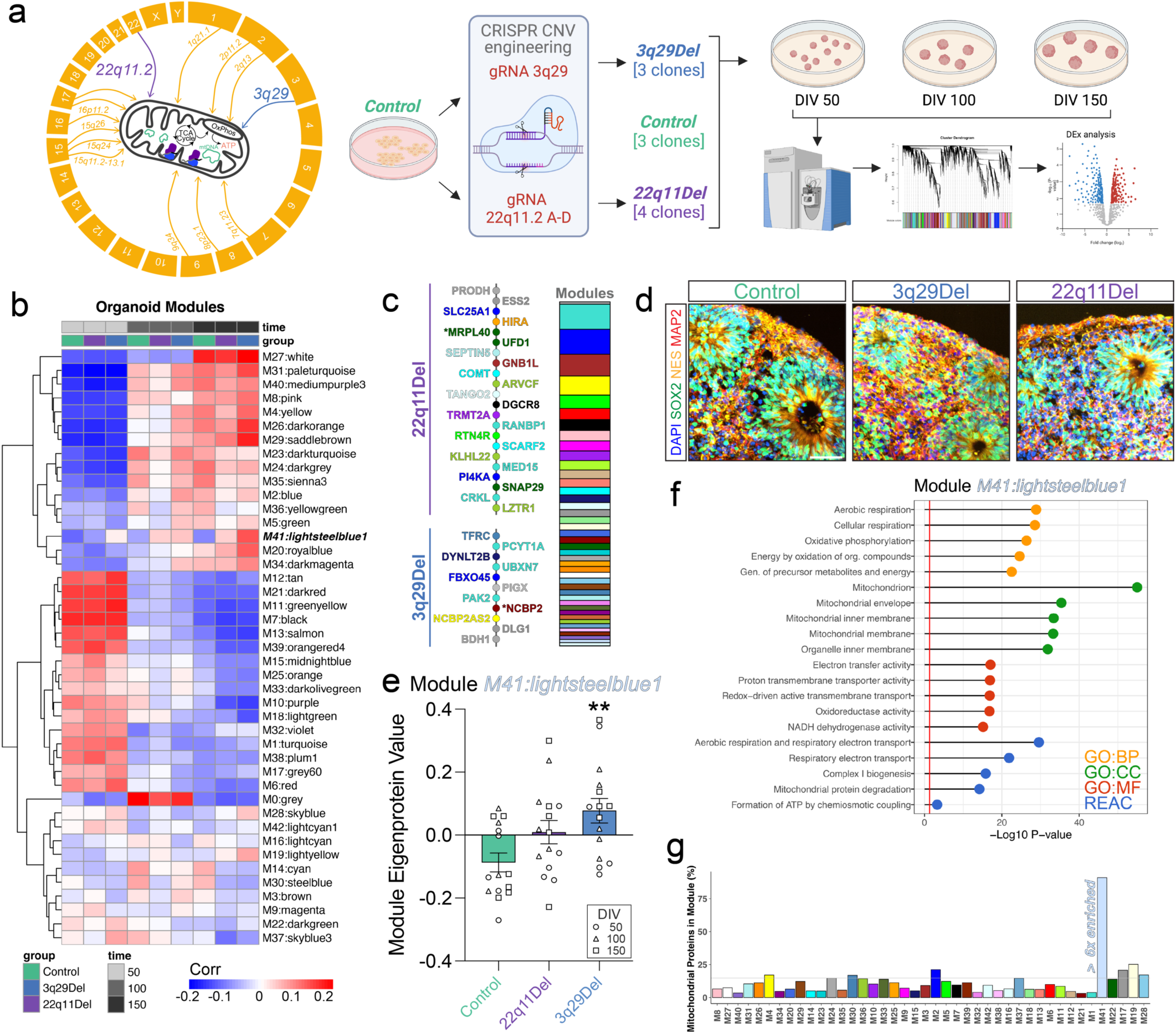
Mitochondrial protein module dysregulated in isogenic organoids. (a) Illustration of experimental design. Interactome networks of multiple NDD CNVs converge on mitochondrial protein networks. Two NDD CNVS, 3q29Del and 22q11Del, were engineered into a control iPS cell line from a neurotypical donor. Clonal lines were differentiated to cortical organoids for 50-150 days and collected for quantitative proteome analyses. (b) Module-trait relationship heatmap. (c) Analyzed proteins encoded by 22q11Del (top) and 3q29Del (bottom) genes are shown in chromosomal order (centromere→telomere) with assigned module colors. Relative module sizes are depicted in the stacked bar plot. CNV proteins found to be module hub genes (kME>0.9) are indicated with asterisks. (d) Example images of dorsal cortical organoids from isogenic cohort at DIV 50 (scale = 50um). (e) Module Eigenprotein value boxplot by genotype of M41 (mean +/- SEM, one-way ANOVA, *post-hoc* Control vs 3q29Del *P = 0.0075*\*\*, Control vs. 22q11Del *P = 0.159*). All time points included for each genotype. (f) Pathway analysis of M41 proteins shows enrichment for mitochondrial ontologies. (g) Analysis of mitochondrial proteins in each module. Expected number (indicated by grey line) is based on the percent of mitochondrial (Mitocarta3.0) proteins in entire dataset (1216/8194 = 14.84%). M41 module had X number of proteins.

**Figure 3.**
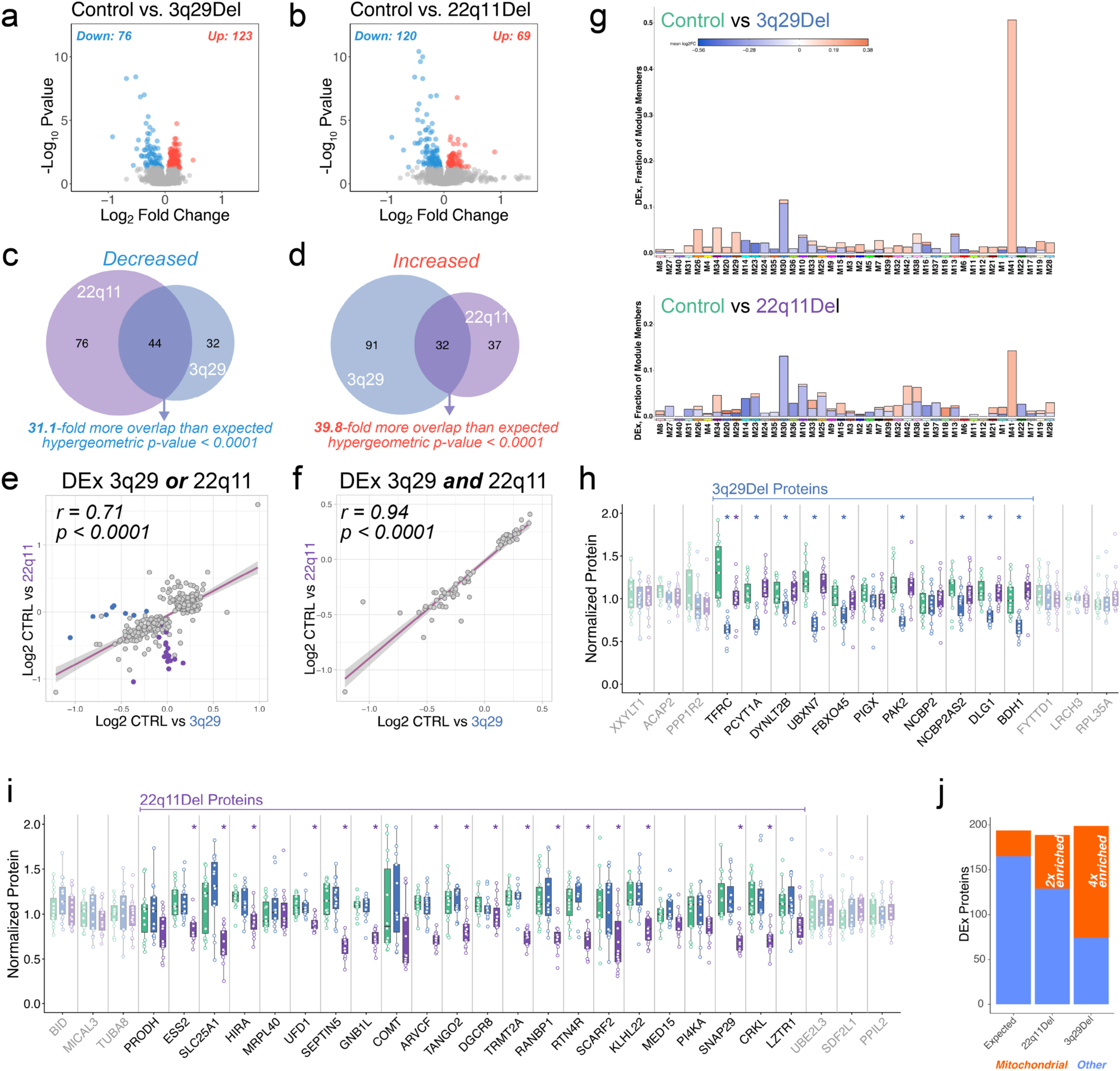
3q29Del and 22q11Del similarly disrupt the developing organoid proteome. (a) Volcano plot of differentially expressed (DEx, P<0.05) proteins in 3q29Del organoids (N=15/genotype). (b) Volcano plot of DEx proteins in 22q11Del organoids (N=15/genotype). (c) Venn diagrams with hypergeometric test to determine degree of overlap for significantly decreased proteins in 3q29Del and 22q11Del. (d) Venn diagrams with hypergeometric test to determine degree of overlap for significantly increased proteins in 3q29Del and 22q11Del. (e) Spearman correlation of Log_2_-fold change of all proteins that were DEx in either 3q29Del or 22q11Del. (f) Spearman correlation of Log_2_-fold change of all proteins that were DEx in 3q29Del and 22q11Del. (g) Portion of each WGCNA module found to be DEx in 3q29Del (top) and 22q11Del (bottom). (h) Boxplots of protein expression for 3q29Del interval and adjacent (grey) proteins. 9/11 analyzed proteins were DEx. The 3q29 protein TFRC was also DEx in 22q11Del. (i) Boxplots of protein expression for 22q11Del interval and adjacent (grey) proteins. 16/22 analyzed proteins were DEx. No 22q11 proteins were found to be DEx in 3q29Del samples. (j) Enrichment analysis of mitochondrial (Mitocarta3.0) proteins. Expected number is based on mean number of DEx proteins in 3q29Del and 22q11Del and percent of proteome that is mitochondrial.

While most modules (29/43) significantly varied by sample time point (two-way ANOVA, *P < 0.05*), only a single module displayed altered eigengene value by genotype (Fig. 2b,e, *M41:lightsteelblue1*, *P = 0.0101*\*, *post-hoc* Control vs 3q29Del *P = 0.0075*\*\*, Control vs. 22q11Del *P = 0.159*). Pathway analysis of proteins assigned to M41 (77 proteins) revealed that it was strongly enriched for mitochondrial ontologies including GO:BP *Oxidative phosphorylation*, GO:CC *Mitochondrion*, and REACTOME *Complex I biogenesis* (Fig 2f). We then assessed mitochondrial protein enrichment across the entire network by comparing the proportion of Mitocarta3.0 proteins [59] found in each module with the expected number if they were evenly distributed across the network. We found that M41 had more than six times as many mitochondrial proteins as would be expected (Fig. 2g). No 3q29 or 22q11 locus proteins were assigned to M41. Module M41 proteins were coordinately increased in 22q11 and 3q29 across all time points compared to controls (most markedly for 3q29 (Fig. 2e) Together, these data suggest that NDD CNVs may lead to dysregulation of mitochondrial proteins in human cortical organoid cultures.

### 3q29Del and 22q11Del similarly disrupt the developing cortical organoid proteome

WGCNA and further exploratory analyses (Supp. Fig. 2) indicated that while many proteins showed dynamic regulation across organoid development, those that were altered by genotype remained largely consistent at each time point. Therefore, we performed differential expression (DEx) analysis by genotype using one-way ANOVA. This analysis revealed 199 DEx proteins (123 increased, 76 decreased) in 3q29Del (compared to Control) and 189 DEx proteins (69 increased, 120 decreased) in 22q11Del (Fig. 3a-b). Nearly half of M41 proteins were significantly increased in 3q29Del along with approximately 15% of M41 proteins in 22q11Del (Fig. 3g). Overall, 3q29Del and 22q11Del produced more dysregulation of mitochondrial proteins than would be expected (Fig. 3j).

To broadly assess the similarity of the impact of 3q29Del and 22q11Del on the developing human cortical organoid proteome, we took two approaches. First, we determined the degree of overlap of increased and decreased DEx proteins in each genotype. Remarkably, among the 76 proteins that were decreased in 3q29Del, a majority (44) were also found be to reduced in 22q11Del, which is >31x more than would be expected by chance (Fig. 3c, hypergeometric *P < 0.0001*\*\*\*\*). Similarly, among the 69 proteins found to be increased in 22q11Del, 32 were also increased in 3q29Del, far more than would be expected (Fig. 3d, hypergeometric *P < 0.0001*\*\*\*\*). Furthermore, we found that the log_2_ fold change of proteins that were found to be DEx in either 3q29Del or 22q11Del organoid compared to Controls showed significant correlation, with many apparent outliers being proteins encoded in these deletion loci (Fig. 3e, Spearman r=0.71, *P < 0.0001*\*\*\*\*). We next pooled the up- and down-regulated protein sets and examined their overlap. Restricting the analysis to the 76 proteins DEx in both genotypes further strengthened the correlation (Fig. 3f, Spearman r=0.94, *P < 0.0001*\*\*\*\*). Lastly, we found that most CNV proteins detected were significantly reduced (Fig. 3h, 9/11 3q29, 3i, 16/22 22q11Del). These results indicate that 3q29Del and 22q11Del have remarkably similar effects on the developing human cortical organoid proteome.

These developing cortical organoid proteome alterations were found to be stable across samples (Fig. 4a), differentiations, and proteomic analysis techniques at two institutions in this collaborative study. In a subsequent proteomic experiment utilizing data-independent acquisition (DIA) described below (Fig. 5), despite smaller cohorts (N=4 vs 15/genotype), 10 of the 76 3q29Del and 22q11Del DEx proteins (Fig. 3f) were again found to be similarly altered in both 3q29Del and 22q11Del (Supp. Fig. 3). We found strong correlations between the fold changes of DEx proteins in the original TMT experiment and in DIA analysis (Supp. Fig. 3, Spearman r = 0.94 for 3q29Del and 0.87 for 22q11Del) supporting the robustness of these results.

**Figure 4.**
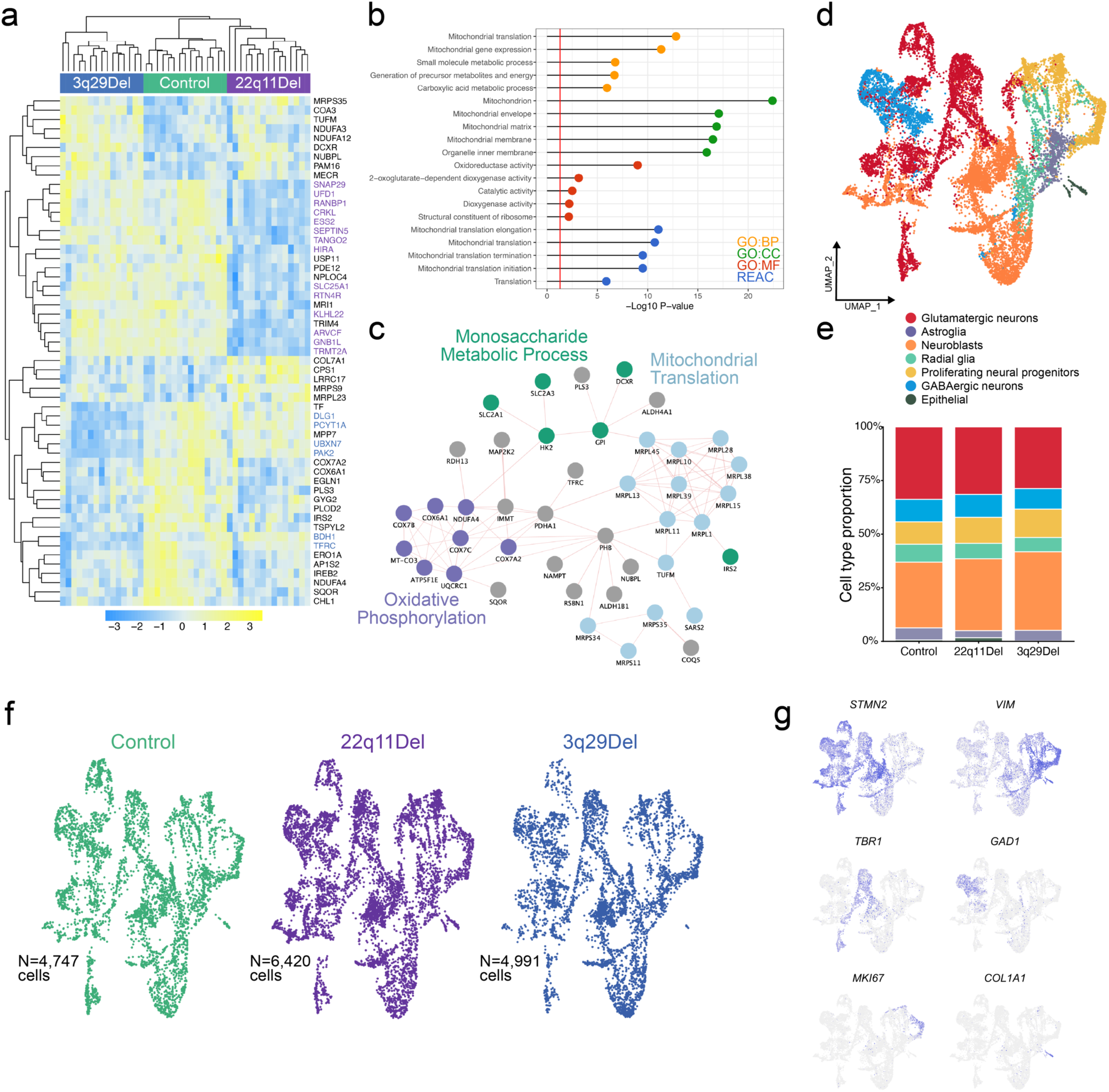
Dysregulated proteins and molecular pathways but stable cell type proportions in 3q29Del and 22q11Del. (a) Heatmap of all DEx proteins at P<0.001 shows distinct clusters of CNV proteins (3q29Del in blue, 22q11Del in purple) and protein sets altered in one or both genotypes. (b) Pathway analysis of 76 common DEx proteins in 3q29Del and 22q11Del with top pathways from each database (GO:BP, GO:CC, GO:MF, REAC) plotted. Red line indicates threshold for statistical significance. (c) More than half (42) of the 76 proteins DEx in 3q29Del and 22q11Del are known to be involved in protein-protein interactions (indicated by pink edges). (d) UMAP of DIV 50 cells from Control, 22q11Del, and 3q29Del. (e) Cell type proportions were not altered by genotype. (f) UMAP split by genotype. (g) Feature plots of key cell type marker genes.

**Figure 5.**
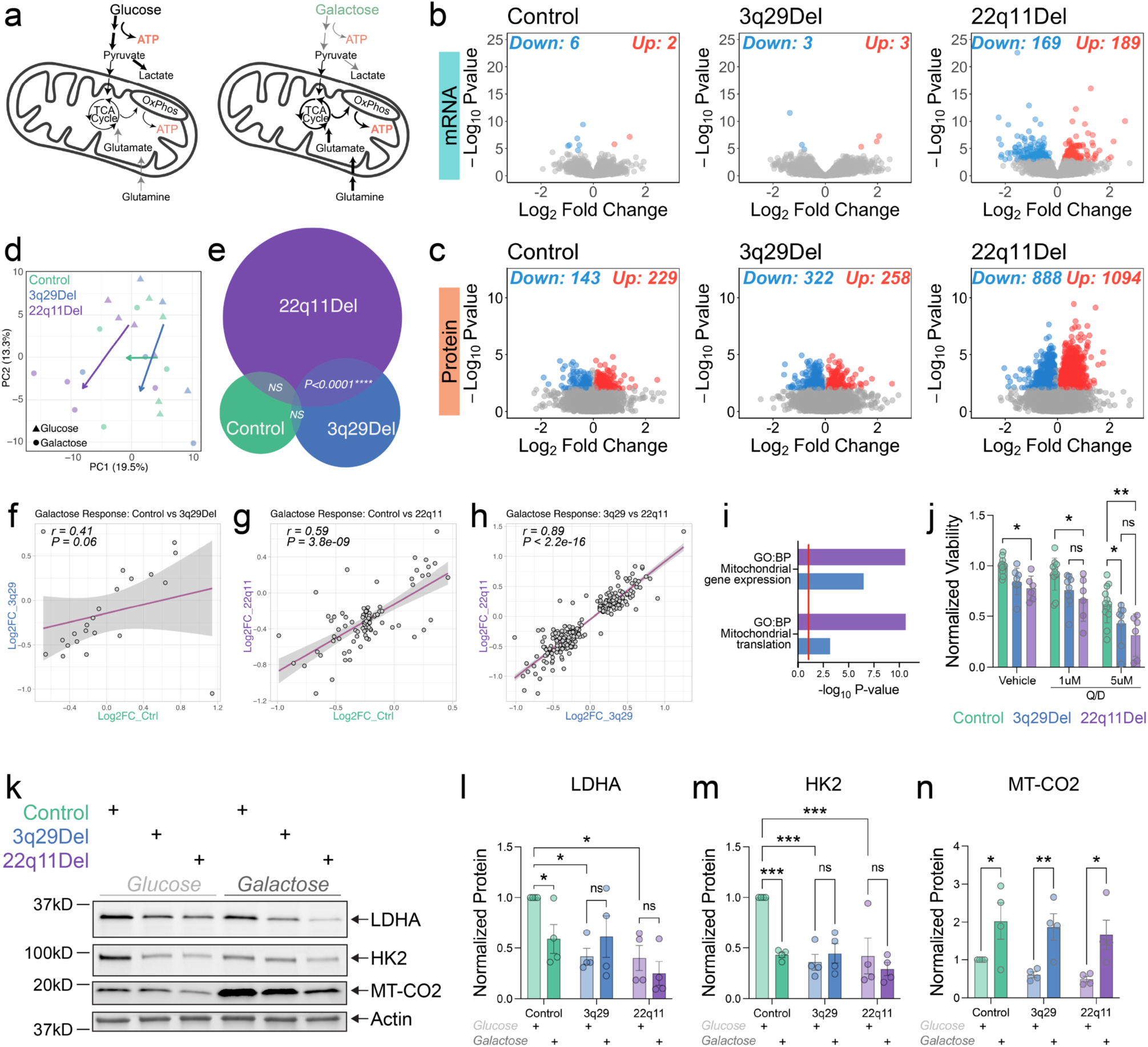
3q29Del and 22q11Del organoids show a similar response to metabolic challenge. (a) Depiction of cellular response to galactose challenge focused on mitochondrial pathway adaptations. (b) Galactose challenge had very little impact on gene expression in Control and 3q29Del organoids but results in more than 350 DEx genes in 22q11Del (*P_adj._<0.05*). (c) Proteomic analysis also showed that 22q11Del organoids had the largest magnitude of DEx protein response (*Q<0.05, Log2FC>0.1*). (d) Principal component analysis of each sample with genotype mean shift from glucose to galactose indicated by arrows. (e) Euler plot of DEx protein overlap by genotype. 3q29Del and 22q11Del showed significant similarity (Overlapping *P-value=7.3e-41*). (f) Spearman correlation of galactose DEx proteins in Control and 3q29Del. (g) Spearman correlation of galactose DEx proteins in Control and 22q11Del. (h) Spearman correlation of galactose DEx proteins in 3q29Del and 22q11Del. (i) Barplot of GO:Biological Process terms enriched among dysregulated proteins in galactose medium (each genotype vs. control, red line indicates significance threshold). (j) Viability of 3q29Del and 22q11Del human neural progenitor cells in vehicle, 1uM, 5uM, of mitochondrial translation inhibitor quinupristin/dalfopristin (N=6-12, two-way ANOVA, main effect of treatment F_(2,69)_=43.14, \*\*\*\**P<0.0001* and genotype F_(2,69)_=16.84, *\*\*\*\*P<0.0001*; post-hoc Tukey comparison \**P<0.05, **P<0.01, ***P<0.001*). (k) Example Western blots from 2-month organoids in glucose medium or challenged with 6-days of galactose medium. (l) Quantification of LDHA Western blots normalized to Actin. N=4/genotype. Two-way ANOVA, main effect of genotype F_(2,18)_=6.8, *P=0.0063*. Asterisks indicate post-hoc Tukey comparisons. (m) Quantification of HK2 Western blots normalized to Actin. N=4/genotype. Two-way ANOVA, main effects of genotype F_(2,18)_=8.9, *P=0.0020*, treatment F_(1,18)_=7.1, *P=0.0154*, and interaction F_(2,18)_=6.4, *P=0.0077*. Asterisks indicate post-hoc Tukey comparisons. (n) Quantification of MT-CO2 Western blots normalized to Actin. N=4/genotype. Two-way ANOVA, main effect of treatment F_(1,18)_=22.84, *P=0.0002*. Asterisks indicate post-hoc Tukey comparisons.

### DEx proteins are enriched for mitochondrial pathways

To initially determine what biological processes may be altered by 3q29Del and 22q11Del in human cortical organoids, we performed pathway analysis on the 76 commonly DEx proteins. These results (Fig. 4b, plotted in order by descending -log_10_ P-value) reinforced previous indications of mitochondrial convergence and revealed a new signal for altered mitochondrial translation. In fact, more than half of these 76 proteins were found to engage in protein-protein interactions (Fig. 4c), suggesting that these proteomic findings implicate a discrete and biologically coherent set of proteins.

### Stable cell-type proportions in 3q29Del and 22q11Del organoids

A possible driver of bulk proteomic changes in 3q29Del and 22q11Del organoids could be altered cell type proportions. To address this possibility, we profiled the transcriptomes of more than 16,000 organoid cells sampled from multiple dissociated organoids of each genotype using single-cell RNAseq (Fig. 4d). More than 25% of cells from all three genotypes were identified as glutamatergic excitatory neurons (*TBR1, BCL11B*) as expected. In addition, we found large proportions of neuroblasts (*DCX*, *NES*), proliferating neural progenitors (*MKI67, VIM*), and radial glia (*PAX6*, *PTPRZ1*) along with a cluster of *GAD1* expressing GABAergic neurons.

We found no statistically significant differences among genotypes in cell type proportions (Fig. 4e, f) as determined by the *propeller* function of R package *speckle* [63] (ANOVA P-values > 0.27). A recent study of 70-150 day organoids reported a delay in excitatory neurogenesis in 22q11Del [69]. Our scRNA-seq data do not entirely rule out this possibility but rather indicate that altered cell type proportion is an unlikely driver of the proteomic changes described in this study.

### 3q29Del and 22q11Del organoids show similar response to metabolic challenge

Differential protein expression analysis results led us to hypothesize that 3q29Del and 22q11Del disrupt mitochondrial function through altered mitochondrial translation. To test this hypothesis, we metabolically challenged Control, 3q29Del, and 22q11Del organoids with equimolar galactose medium, which blunts glycolysis and can reveal mitochondrial deficits that are concealed in glucose medium conditions (Fig. 5a) [70, 71]. Our previous work showed that control neural cells increase oxygen consumption rate in this glucose to galactose challenge paradigm, yet this response is ameliorated in 3q29Del [43]. We reasoned that shifting cells from glycolysis to oxidative phosphorylation (OxPhos) would serve as a metabolic challenge to neural cells with mitochondrial translation deficiencies.

After 5 weeks of differentiation in standard glucose media, we transferred a set of organoids to galactose medium for one week and collected samples for transcriptomic and proteomic analysis. Bulk RNA-seq data showed very few changes in Control and 3q29Del organoids, but surprisingly more than 350 transcripts were significantly altered in 22q11Del organoids (Fig. 5b, glucose vs. galactose, *P_adj._ < 0.05*). Proteomic changes were obvious in all three genotype groups, but 22q11Del displayed by far the most disruption with more than 1,900 proteins significantly altered (Fig. 5c, *q<0.05*, |Log_2_FC| > 0.1).

To assess the similarity of responses to the metabolic challenge in 3q29Del and 22q11Del, we performed overlap, correlation, and principal component (PC) analyses. PC analysis of the proteomic shift from glucose to galactose indicated qualitatively similar effects in both 3q29Del and 22q11Del (Fig. 5d). There was no significant overlap in proteomic response to galactose between Control and 3q29Del organoids (Fig. 5e) and no significant correlation (Fig. 5f). Control and 22q11Del also showed no significant overlap (Fig. 5e) but the overlapping DEx proteins were significantly correlated in responses to galactose shift (Fig. 5g, Spearman r = 0.59, *P = 3.8e-9*\*\*\*\*). However, 3q29Del and 22q11Del were found to have significant overlap (Fig. 5e, 2.1x overlap, *P < 0.0001*\*\*\*\*) and remarkable correlation (Fig. 5h, r = 0.89, *P < 2.2e-16\*\*\*\**). Pathway analysis revealed that GO:Biological Process terms *Mitochondrial translation* and *Mitochondrial gene expression* were significantly altered in both 3q29Del and 22q11Del organoids by the galactose metabolic challenge (Fig. 5i). These results suggest that developing cortical cells carrying 3q29Del and 22q11Del mount similar responses to metabolic challenge that differ from intact cell responses.

In a subsequent differentiation, we again performed the galactose challenge and collected cortical organoid protein samples for targeted immunoblotting experiments. Galactose medium is expected to target enzymes of the central carbon metabolism reducing cellular reliance on glycolysis and increase oxidative phosphorylation. Indeed, the key glycolysis enzymes LDHA and HK2 were identified as DEx in both 3q29Del and 22q11Del. Immunoblotting showed that the galactose treated Control organoids had diminished LDHA and HK2. However, 3q29Del and 22q11Del organoids displayed lower LDHA and HK2 protein levels at baseline in glucose medium and the galactose challenge did not significantly alter LDHA or HK2 expression (Fig. 5k-m). We predicted that the galactose challenge would lead to increased expression of mtDNA-encoded proteins, which may be altered in 3q29Del and 22q11Del. As we expected, the galactose challenge did in fact significantly increase MT-CO2 protein in Control organoids, yet 3q29Del and 22q11Del organoids also showed this response and there was no significant difference in immunoblot densitometry by genotype (Fig. 5k, n).

### Mitochondrial translation inhibition reduces viability in 3q29Del and 22q11Del neural progenitor cells

To further test the hypothesis that 3q29Del and 22q11Del impair mitochondrial translation in human neural cells, we treated neural progenitor cell (NPC) cultures from Control, 3q29Del, and 22q11Del cell lines with the antibiotic cocktail quinupristin/dalfopristin (Q/D), which has been shown to potently inhibit mitochondrial translation [65]. We found that Q/D treatment diminished the viability of all NPC lines in the resazurin metabolism assay, but 3q29Del and 22q11Del NPCs were impacted significantly more than Control cells (Fig. 5j).

Together, these data support a model where haploinsufficiency of distinct mitochondrial interactome nodes convergently disrupts core functions including mitochondrial translation.

## Discussion

A central unresolved question in NDD genetics is how genetically distinct high-effect variants produce convergent clinical phenotypes. Our findings provide a mechanistic insight to address this issue. Rather than acting through entirely unique pathways driven by discrete genetic variants, we show that recurrent CNVs and SCZ risk genes converge on a shared cellular vulnerability centered on mitochondrial translation. Network analysis demonstrates that risk genes are frequently positioned within a highly connected mitochondrial protein interactome. Critically, we demonstrate that two highly penetrant CNVs with substantial phenotypic overlap, 3q29Del and 22q11Del [7, 22, 72], produce strikingly similar proteomic alterations in isogenic human cortical organoids and converge on shared molecular pathology, producing highly concordant proteomic, metabolic, and functional deficits in a model of developing human cortex. Taken together, these data indicate that NDD risk variants do not require the disruption of identical genes to produce similar disease risk; instead, they converge on a biological process, mitochondrial translation, that impacts metabolic adaptability in neuronal development. The interactome analyses provide a conceptual framework for this convergence. Although mitochondrial genes are not strongly enriched at the level of gene annotation in psychiatric genetics, CNV-encoded and schizophrenia risk proteins occupy highly connected positions within a mitochondrial proximity interaction network (Fig. 1). This indicates that disease mechanisms need not arise from direct disruption of canonical mitochondrial genes. Rather, high-risk variants may influence mitochondria indirectly through network propagation. Interaction partners of risk genes were preferentially enriched for mitochondrial gene expression and translation pathways, suggesting that this subsystem represents a constrained functional node within the broader proteome. These observations position mitochondria as a central integrative hub through which genetically diverse psychiatric risk variants converge to impose shared cellular vulnerability.

We directly tested this network convergence hypothesis in an isogenic human neurodevelopmental system. Comparison of 3q29Del and 22q11Del cortical organoids showed that genotype-dependent effects converged on a mitochondrial-enriched module that did not contain CNV-encoded proteins (Fig. 2e-g, Fig. 3h-i). Both CNVs produced highly overlapping proteomic alterations with shared dysregulation of mitochondrial gene expression and translation pathways (Figs. 3f, Fig. 4b-c). While mitochondrial dysfunction has been implicated in both 3q29Del and 22q11Del by our group and others [43, 46, 48, 73], our findings support a downstream network of effects of oligogenic haploinsufficiency. Additionally, convergence represents a functional liability that we unmasked by shifting metabolic demand. When challenged to organoids to prioritize OxPhos, both CNVs displayed a remarkably similar proteomic response and blunted responsiveness to key glycolytic enzymes (Fig. 5h, k-m). These data support the model of metabolic inflexibility that we described previously for 3q29Del [43]. Consistent with this interpretation, pharmacologic inhibition of mitochondrial translation disproportionately impacted 3q29Del and 22q11Del NPC viability (Fig. 5j). Together, these findings provide orthogonal functional evidence that mitochondrial translation capacity is a critical limiting factor in these high-risk CNVs. Consistent with this interpretation, independent evidence from human disease tissue points to a similar vulnerability. We previously observed reduced expression of mitochondrial ribosomal subunit transcripts in pyramidal and parvalbumin neurons from post-mortem schizophrenia prefrontal cortex [47], suggesting that impaired mitochondrial translation may extend beyond CNV models to idiopathic psychiatric disorders.

Future work should explicitly investigate how synaptic and mitochondrial dysfunction converge in 3q29Del and 22q11Del. Although these neuronal domains are often treated as independent in the literature, mitochondria are integral to synaptic function, comprising up to one-third of synaptic volume and scaling with synaptic activity across neural circuits [74, 75]. Consistent with this functional interdependence, our bioinformatic analyses indicate that 22q11Del robustly impacts both synaptic and mitochondrial gene networks. Specifically, 19 of the 20 NDD CNVs analyzed contain SynGO-annotated synaptic genes, while 18 of 20 contain Mitocarta 3.0-annotated mitochondrial genes (Fig. 1a). Multiple studies have reported synaptic impairments in mouse models of 22q11Del [76], although it remains unclear whether these phenotypes arise directly from dysregulation of synaptic proteins or indirectly through mitochondrial dysfunction. Dysregulated calcium signaling, a process tightly linked to both mitochondrial function and synaptic plasticity, has been implicated in these models [77]. Notably, haploinsufficiency of the mitochondrial ribosomal protein MRPL40 alters mitochondrial calcium handling, resulting in deficits in short-term synaptic plasticity, working memory, and prepulse inhibition [68, 78]. Complementing these findings, impaired neuronal activity and calcium signaling have been observed in human iPSC-derived neurons and brain organoids from individuals with 22q11Del [79] Together, these studies motivate further investigation into how 22q11Del genes influence synaptic function through mitochondrial pathways, particularly mitochondrial translation.

In contrast, the mechanisms by which 3q29Del genes converge on synaptic and mitochondrial biology are less well defined. Early studies emphasized postsynaptic signaling complexes, largely due to DLG1, a synaptic scaffolding protein encoded within the 3q29 interval [80]. However, our work demonstrated that *Dlg1* haploinsufficiency alone does not recapitulate the behavioral phenotypes observed in 3q29Del mouse models [81], suggesting that additional genes or combinatorial effects are required. One compelling 3q29-encoded candidate linking synaptic signaling and mitochondrial metabolism is PAK2, a serine/threonine kinase with established roles in synapse function. PAK2 haploinsufficiency produces synaptic and behavioral deficits in mice [82] and has been implicated in glucose uptake and insulin sensitivity [83]. Consistent with these metabolic roles, our recent work demonstrated that PAK2 haploinsufficiency contributes to metabolic inflexibility in a human cellular model of 3q29Del [43]. Collectively, these findings support the existence of proteins encoded within the 3q29 interval that exert convergent effects on synaptic signaling and mitochondrial metabolism. However, whether and how these mechanisms selectively impact mitochondrial translation remains an important unresolved question.

Several limitations define important avenues for future investigation. Although our cortical organoid differentiation approach effectively recapitulates early stages of human corticogenesis, the lack of vascularization, immune integration, and long-range circuit activity likely limits metabolic demand, potentially underestimating the consequences of mitochondrial translation constraints present in more physiologically complex systems [84, 85]. In addition, the preservation of CNV-encoded hub proteins NCBP2 and MRPL40 suggests that cells preferentially maintain key network hub proteins and implicates the presence of compensatory post-transcriptional mechanisms that are not fully resolved by bulk proteomic approaches. Dissecting these regulatory layers will require higher-resolution strategies to determine how such compensatory processes are maintained and where they ultimately fail. Finally, establishing causality and generalizability will require future studies incorporating multi-line validation and gene-specific rescue experiments to definitively map the molecular trajectory from genomic deletion to metabolic failure.

These findings have several implications. First, they suggest that NDD-associated risk variants may converge on mitochondria through interaction network architecture, rather than through canonical mitochondrial genes, impacting mitochondrial translation as a common cellular vulnerability. Importantly, several recent studies from post-mortem tissue and iPSC-based models add further support to the model of transcriptomic convergence across NDD risk variants. A cortical organoid study of iPSCs carrying eight different ASD-associated variants, including 22q11Del, found strong evidence for transcriptomic convergence across disorders, which, in agreement with our findings, were stable across in vitro maturation time points [86]. Moreover, a single-nucleus transcriptomic study of carriers of nine NDD-related CNVs including 22q11Del found strong evidence for dysregulated gene expression converging across variants on mitochondrial energy production [87]. A pair of recent single nucleus transcriptomic and bulk proteomic studies of cortical samples from several dozen individuals with SCZ and neurotypical controls found significant down-regulation of mitochondrial OxPhos genes and proteins [15, 88]. Together, these observations suggest that mitochondrial dysfunction, perhaps driven by inefficient translation represents a shared, systems-level constraint on neuronal development through which genetically diverse risk variants can produce convergent disease phenotypes.

In summary, our results support a model in which NDD-associated CNVs and SCZ risk genes converge on a mitochondrial interaction network enriched for mitochondrial translation and gene expression processes. We show that 3q29Del and 22q11Del produce highly concordant proteomic alterations in developing human cortical organoids and exhibit heightened sensitivity to metabolic and pharmacologic constraints on mitochondrial gene expression. Collectively, these findings strengthen the case for mitochondrial translation as a shared vulnerability across genetically distinct NDD risk variants.

## Acknowledgments

This work was supported by a Brain & Behavior Foundation Young Investigator Award (RHP), seed funding from the Emory University Research Committee (RHP), the Emory University School of Medicine (*I^3^ Nexus Award* JGM, JFC, ED, GJB, *I^3^ SOM-VA Award* RHP, ED, GJB), the US National Institute of Mental Health (K01 MH133970 to RHP, R36 MH136806 to MIR, R01MH125956 to SAS, R01MH117315 to JFC, ED, R01 MH110701 to JGM), the US National Institute of Environmental Health Sciences (R01 ES034796), and the Fralin Biomedical Research Institute at Virginia Tech Carilion.

This study was supported in part by the Emory Integrated Proteomics Core (RRID:SCR_023530), which is subsidized by the Emory University School of Medicine and is one of the Emory Integrated Core Facilities. Additional support was provided by the Georgia Clinical & Translational Science Alliance of the National Institutes of Health under Award Number UL1TR002378.

Author ED is an employee of the US government. The content is solely the responsibility of the authors and does not necessarily reflect the position or policy of the National Institutes of Health, the Department of Veterans Affairs, or the US government. ED has received research support for work unrelated to this project from the Department of Defense.

## Supplemental Figures

**Supplemental Figure 1.**
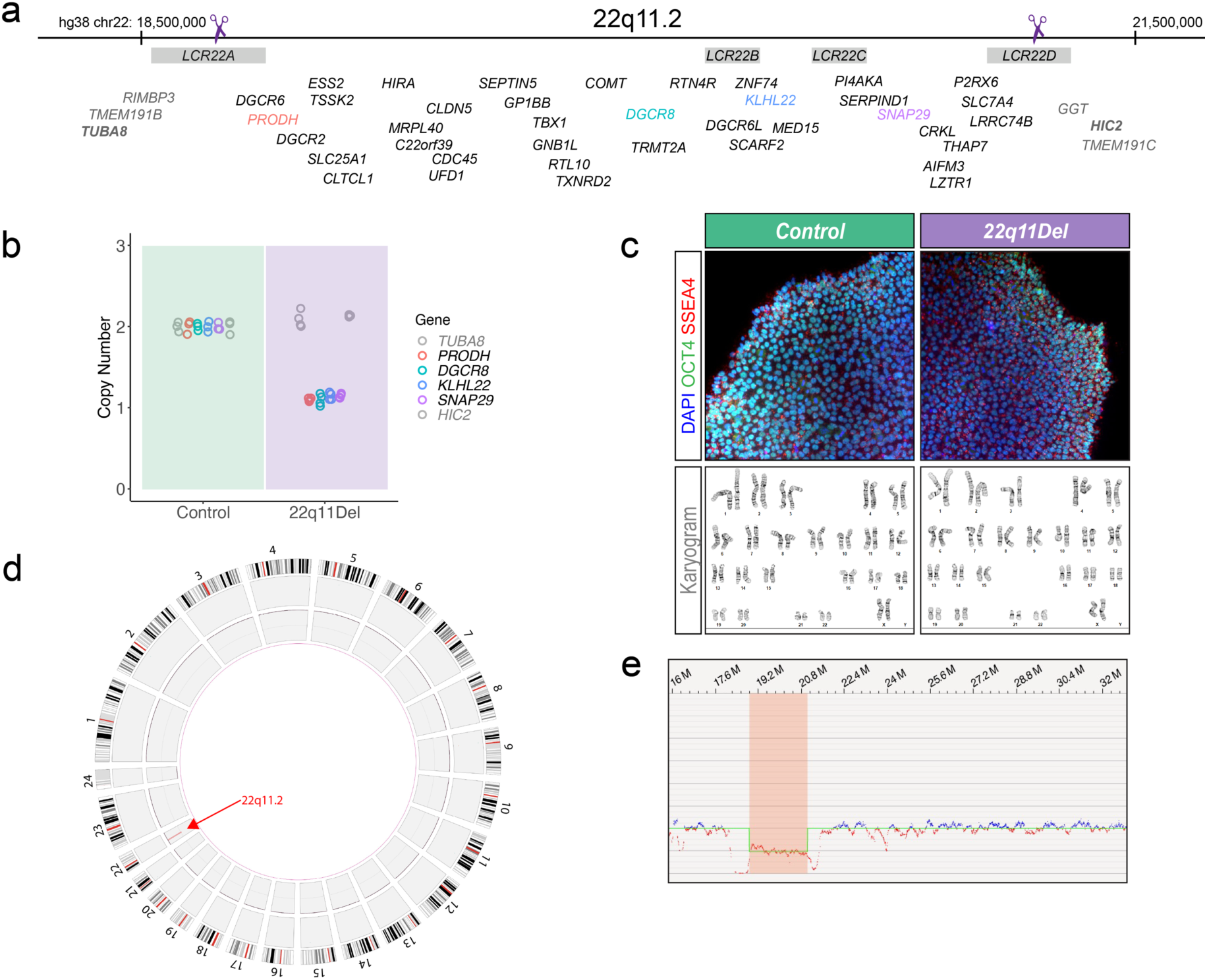
Isogenic 22q11.2 deletion iPSC lines. (a) Illustration of 22q11.2 deletion locus. A single gRNA targeted repeated sequence in LCR-A and LCR-D. (b) TaqMan copy number assay results for isogenic 22q11Del clones. Bolded grey genes flanking the deletion interval and one color-highlighted gene within each LCR segment (a) were probed. (c) Immunofluorescence for pluripotency markers OCT4 and SSEA4 showed isogenic 22q11Del cells maintained typical iPSC morphology. (d) Circos plot of genome-wide optical mapping data shows the 22q11Del is the only new structural variant. (e) Genome-wide optical mapping traces show a deletion at 22q11.2.

**Supplemental Figure 2.**
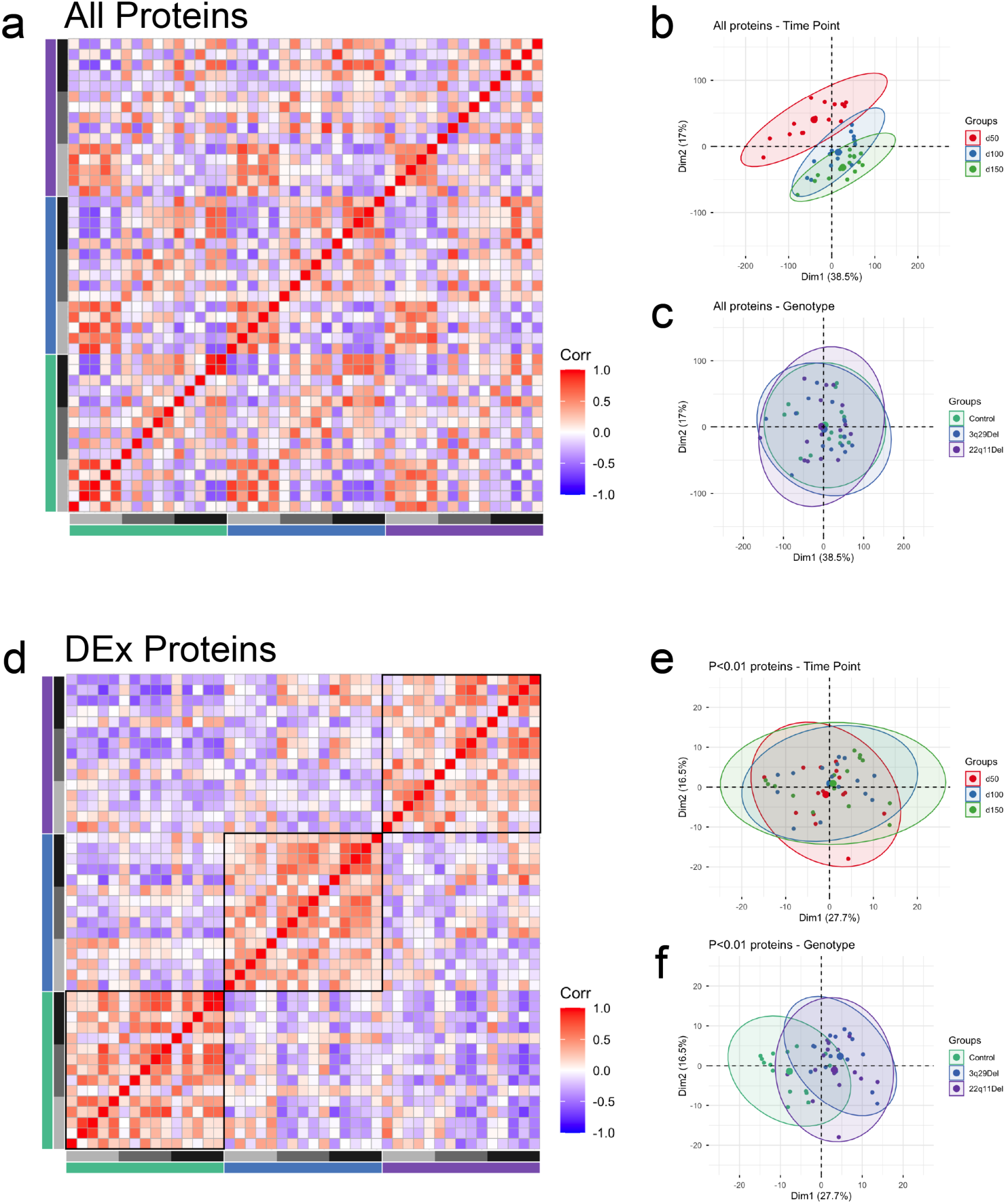
TMT proteomic analysis. (a) Correlation matrix of all samples and all proteins. (b) Principal component analysis (PCA) with samples labeled by time point indicates d50 samples are largely distinct from d100 and d150 regardless of genotype. (c) PCA of all proteins with samples labeled by genotype shows no clear difference by genotype group. (d) Correlation matrix of all samples for all DEx proteins (P<0.01) shows clear clustering by genotype with no discernable time point clustering. (e) PCA of DEx proteins shows no clear distinction by time point. (f) PCA of DEx proteins labeled by genotype shows strong overlap of 3q29Del and 22q11Del that is somewhat distinct from Control.

**Supplemental Figure 3.**
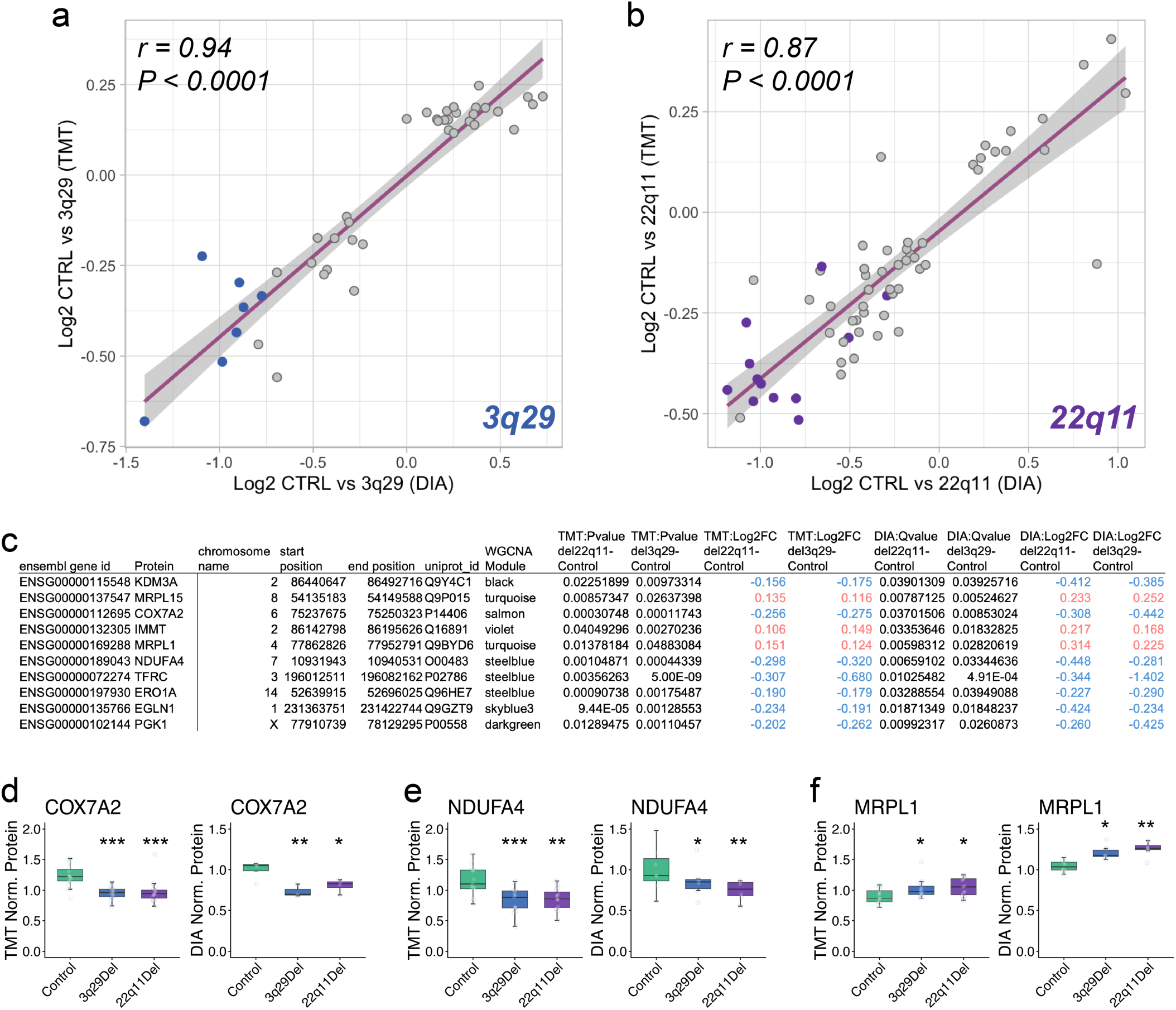
Comparison of TMT and DIA proteomic data sets. (a) Plot of Pearson correlation of Log2FC of proteins significantly DEx in 3q29Del in TMT and DIA experiments (DIA data at DIV50, glucose medium only). Proteins from 3q29Del locus highlighted in blue. (b) Plot of Pearson correlation of Log2FC of proteins significantly DEx in 22q11Del in TMT and DIA experiments. Proteins from 22q11Del locus highlighted in purple. (c) Table of the 10 proteins that were DEx in both 3q29Del and 22q11Del compared to control in both datasets. (d-f) Example box plots of proteins from list of 10 in table c (*P<0.05; **P<0.01, ***P<0.001).

---

**Supplemental Table 1.**

| **Cell Line** | **Clone** | **Genotype** | **Sex** | **Source** | **Experiments** |
| --- | --- | --- | --- | --- | --- |
| 1003-0031 | C19 | Control | Female | Isogenic | TMT, DIA, Western blot, Resazurin |
| 1003-0031 | C27 | Control | Female | Isogenic | TMT, DIA, Western blot, Resazurin |
| 1003-0031 | C33 | Control | Female | Isogenic | TMT, DIA, Western blot, Resazurin |
| 1003-0031 | 3G2b | 3q29Del | Female | Isogenic | TMT, DIA, Western blot, Resazurin |
| 1003-0031 | 3H2 | 3q29Del | Female | Isogenic | TMT, DIA, Western blot |
| 1003-0031 | 11D7 | 3q29Del | Female | Isogenic | TMT, DIA, Western blot, Resazurin |
| 1003-0031 | 1F3 | 22q11Del | Female | Isogenic | TMT, DIA, Western blot, Resazurin |
| 1003-0031 | 3D8 | 22q11Del | Female | Isogenic | TMT, DIA, Western blot |
| 1003-0031 | 6F4 | 22q11Del | Female | Isogenic | TMT, DIA, Resazurin |
| 1003-0031 | 10C8 | 22q11Del | Female | Isogenic | TMT, DIA, Western blot |
| 22q#3 | C1 | 22q11Del | Female | Study | Resazurin |
| 22q#4 | C6 | Control | Female | Study | Resazurin |
| 4258-2046 | C17 | Control | Male | Study | Resazurin |
| 4258-2046 | C19 | Control | Male | Study | Resazurin |
| 4258-2046 | 1C7 | 3q29Del | Male | Isogenic | Resazurin |
| 4169-1001 | C14 | 3q29Del | Male | Study | Resazurin |

